# Patterns of microbial colonization of human bone from surface-decomposed remains

**DOI:** 10.1101/664482

**Authors:** Alexandra L. Emmons, Amy Z. Mundorff, Sarah W. Keenan, Jonathan Davoren, Janna Andronowski, David O. Carter, Jennifer M. DeBruyn

## Abstract

Microbial colonization of bone is an important mechanism of post-mortem skeletal degradation. However, the types and distributions of bone and tooth colonizing microbes are not well characterized. It is unknown if microbial communities vary in abundance or composition between bone element types, which could help explain patterns of human DNA preservation. The goals of the present study were to (1) identify the types of microbes capable of colonizing different human bone types and (2) relate microbial abundances, diversity, and community composition to bone type and human DNA preservation. DNA extracts from 165 bone and tooth samples from three skeletonized individuals were assessed for bacterial loading and microbial community composition and structure. Random forest models were applied to predict operational taxonomic units (OTUs) associated with human DNA concentration. Dominant bacterial bone colonizers were from the phyla Proteobacteria (36%), Actinobacteria (23%), Firmicutes (13%), Bacteroidetes (12%), and Planctomycetes (4.4%). Eukaryotic bone colonizers were from Ascomycota (40%), Apicomplexa (21%), Annelida (19%), Basidiomycota (17%), and Ciliophora (14%). Bacterial loading was not a significant predictor of human DNA concentration in two out of three individuals. Random forest models were minimally successful in identifying microbes related to patterns of DNA preservation, complicated by high variability in community structure between individuals and body regions. This work expands on our understanding of the types of microbes capable of colonizing human bone and contributing to human skeletal DNA degradation.

## Introduction

Skeletonization is the final stage of human decomposition, exposing bone to the surrounding environment [1]. Once the body has progressed to a skeletonized state, teeth and bone become the only materials that can be used for DNA identification. However, while bone is more recalcitrant than soft tissue, it is not stable; it continues to decay over time. With death, bone undergoes decomposition and diagenesis, the postmortem alteration of bone by chemical, physical, and biological factors that result in modification of the original bone material [2]. Time alone is not a good indicator of skeletal DNA preservation [3]. Instead, bone diagenesis and DNA survival is highly dependent on the depositional environment, including microbial activity [4,5], just as the decomposition of all organic resources are influenced by the decomposer community and physicochemical environment [6].

Bone decay mechanistically proceeds via chemical and/or microbial degradation of the organic and inorganic components of bone [7]. Microbes are capable of colonizing and degrading human bone, and microbial DNA is often co-extracted with human DNA, which interferes with downstream processes [8–10]. The organic component of bone consists of 90-95% type I collagen (primarily made up of glycine, proline, and hydroxyproline), with minor contributions from other non-collagenous proteins (e.g., osteocalcin, osteoponin, and osteonectin) as well as lipids, mucopolysaccharides, and carbohydrates [11]. The inorganic or mineral component is most similar to hydroxyapatite and consists of calcium, phosphate, carbonate, and to varying degrees sodium [2,12,13]. Bone apatite, or bioapatite, can be described as ‘nature’s trashcan,’ as infiltration and substitutions for environmental elements are common [2]. One of the main requirements for lasting preservation via fossilization is a complete shift from bioapatite composition to a more stable mineral phase, such as fluorinated apatite or fluorine- and carbonate-enriched apatite [2,14].

When not in equilibrium with the surrounding environment, dissolution and recrystallization of bioapatite occurs, allowing microorganisms and enzymes access to the organic phase, resulting in degradation. Similarly, if the organic component degrades by either chemical or biological means, bioapatite becomes more vulnerable to environmental fluctuations and dissolution of the lattice structure is more probable due to new voids in the crystal lattice [2,7,12,15–17]. For example, wet environments exhibit increased rates of DNA degradation because water allows for mineral dissolution and increased hydrolytic damage [18]. The interdependence between the mineral and organic phases of bone supports the idea that greater porosity increases the susceptibility of bone to environmental influences [19,20], both chemical and biological.

Though the reservoir for long-term DNA preservation in bone remains unclear, binding of DNA to bioapatite crystallites seems to be crucial for long-term DNA survival [15]; persistence within osteocytes or other remnant tissues (e.g., from the red bone marrow) may also be possible [21,22]. Gross bone preservation and weathering has been shown to be unrelated to DNA preservation or degradation in some cases [19], while in others, indices of gross preservation are better correlated [23,24]. Differences in DNA preservation and degradation by bone type have been observed, though patterns are not consistent between studies (e.g., [9,10,25–28]). Whether this has to do with differences in cortical and cancellous bone composition is debated. More porous elements are thought to have increased bacterial presence [15], but increased presence does not necessarily mean increased degradation, as certain microbial taxa may be better adapted to exploiting skeletal material than others.

In archaeology, microbial degradation of bone has been studied primarily through histological research, focusing on regions of microscopic focal destruction [24,29–32]. However, culture-based research has shown that collagenase-producing bacteria can use mammalian bone as a substrate (e.g., *Alcaligenes pichaudii, Bacillus subtilis, Pseudomonas fluorescens, Clostridium histolyticum)* [33]. Others have shown greater DNA preservation from archaeological sites with bones lacking culturable collagenase producing bacteria [34]. These observations suggest that DNA preservation within a bone may be partially dependent on the amount and/or type of microbes colonizing bones. Genera including *Pseudomonas*, *Xanthomonas*, *Fusarium,* and *Trichonella* have been cultured from bones from diverse archaeological sites [34]. Experimental research has also shown macroscopic destruction phenomena consistent with fungal degraders, specifically the genus *Mucor* [35], while others have cultured genera from the phylum Ascomycota [34]. Research to date has primarily been limited to culture-based methods, and only a small subset of environmental microbes can be cultivated in the laboratory [36]. Only a few studies [37,38] have been conducted since the advent of high throughput sequencing technologies, which permit microbial characterization without cultivation. Thus, there is a gap in knowledge regarding the types of microbes capable of colonizing and degrading human bone.

The purpose of the current study was two-fold: (1) to identify the types of microbes capable of colonizing different human bone types using next generation sequencing, and (2) to relate microbial abundances, diversity, and community composition to bone type and patterns of human DNA preservation. We expected total bacterial gene abundances, as a proxy for overall bacterial presence or loading, to increase with decreasing human DNA quantity and quality. We also expected to see shifts in microbial populations with changes in bone morphological and microstructural properties (i.e., specific element type and cortical content).

## Materials and Methods

In 2009, three male individuals were placed outside on the ground surface to decompose naturally at the Anthropology Research Facility (ARF) at the University of Tennessee, Knoxville (UTK) (S1 Table). The individuals were donated to the donated to the University of Tennessee Forensic Anthropology Center for the W. M. Bass Donated Skeletal Collection. Because no living human subjects were involved in this research and no personally identifiable information was collected, the project was exempt from review by the University of Tennessee Institutional Review Board. The skeletons were mapped and recovered following complete skeletonization (13 to 23 months), and gently washed with a new toothbrush and tap water at the Forensic Anthropology Center (UTK). The same 55 bones and teeth from each individual (total n = 165), which represented all skeletal element types, were selected for sampling (Table 1, S2 Fig). Prior to sampling, the external surface of each bone was cleaned by mechanically removing 1 to 2 mm of the outer surface, followed by chemical cleaning via bleach, ethanol, and sterile water. Bones were sampled using a drill and masonry bit at slow speeds; DNA was extracted from sampled bone powder using a complete demineralization protocol [39]. Bone sampling and DNA extraction and analysis were previously described in detail in Mundorff and Davoren [28]. Human DNA quality and quantity were examined to elucidate patterns of DNA preservation by bone type [28]. These remaining skeletal DNA extracts were used in the present study to assess microbial loading via qPCR and microbial community composition and structure using next generation sequencing of the 16S rRNA and 18S rRNA genes.

**Table 1:**
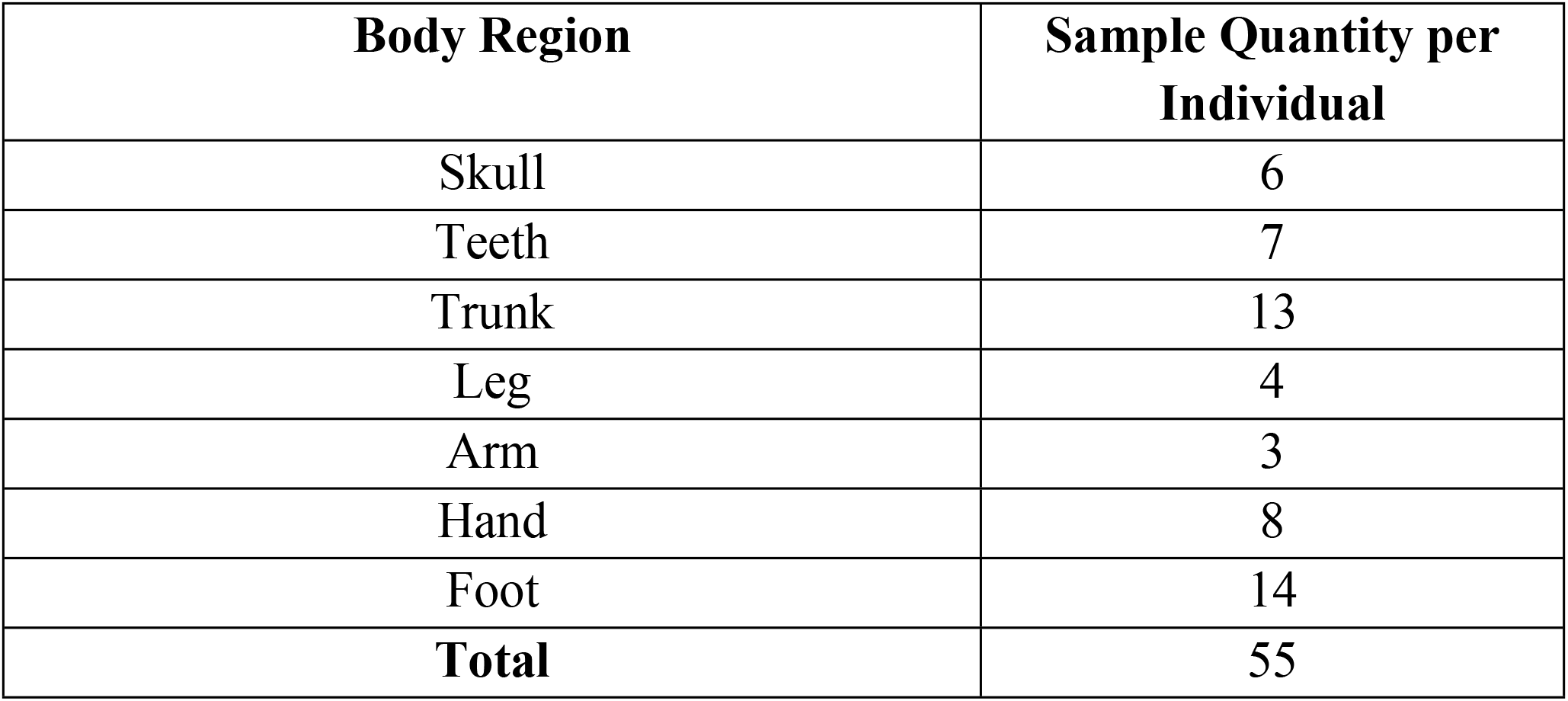
Number of bones sampled by body region for each of the three individuals

| Body Region | Sample Quantity per Individual |
| --- | --- |
| Skull | 6 |
| Teeth | 7 |
| Trunk | 13 |
| Leg | 4 |
| Arm | 3 |
| Hand | 8 |
| Foot | 14 |
| **Total** | **55** |

### Microbial and human DNA quantification

As a proxy for total bacterial abundance and colonization of bone, qPCR was used to quantify 16S rRNA gene abundances [40] using the Femto™ Bacterial DNA Quantification Kit (Zymo Research). Assays were conducted following manufacturer instructions using a BioRad CFX Connect™ Real-Time PCR Detection System. Samples were quantified in triplicate, while standards were quantified in duplicate, and a minimum of three no template controls were included in each 96-well plate. Data are presented as gene copy number per gram of bone powder (gene copies gbp^-1^). Human DNA was quantified using the Quantifiler™ kit from Life Technologies (Qf); methods and data are reported in [28].

### Total DNA quantification

Total DNA was quantified using the Quant-iT™ PicoGreen™ dsDNA Assay Kit (Invitrogen™) using a 200 μL total volume on a 96 well microplate reader. Samples and standards were run in duplicate, with standards ranging from 0 μg mL^-1^ to 1.0 μg mL^-1^. Total DNA concentrations are reported as nanograms per gram of bone powder (ng gbp^-1^).

### Percentage cortical content

Clinical CT scans of each element were acquired using a Siemens Biograph mCT 64 slice scanner. Scans involved helical acquisition using a 0.6 mm slice thickness, 500 mAs, 120 kV, and bone window with kernel B70s. Data were stored on compact discs and transferred to workstations with image processing software (OsiriX 5.6, Geneva, Switzerland). The DNA sampling site on each element was digitally measured using ImageJ (National Institutes of Health). A macro was created to detect and measure the areas of cortical and cancellous bone (mm) on each CT slice where the sampling site appeared. Measurements of cortical width and height and cancellous width and height were taken separately for each cortical and cancellous bone region for all bones. Average cortical and cancellous bone width and height measurements were then computed. Due to issues with scan quality, ten of the original 129 samples were removed from the analysis. Percentages of cortical and cancellous bone were computed from each DNA sampling site for all elements.

Mean percentages of cortical bone composition at each sampling site were divided into seven categories by skeletal element: (1) 80 to 100%, (2) 70 to 79%, (3) 60 to 69%, (4) 50 to 59%, (5) 40 to 49%, (6) 30 to 39%, and (7) 20 to 29%. The first category consists of bones whose sampling sites did not contain any cancellous bone, including the humerus, radius, ulna, femur, and tibia. The second, third, and fourth categories contained only three elements with sampling sites that were composed of over 50% cortical bone. Percentage data were further averaged from each element type across all individuals. The majority of element types revealed consistent measurements between individuals, with standard deviations of 10% or less. Three element types (temporal, occipital, cervical vertebra) exhibited high variability between the three individuals in the relative amount of cortical and cancellous bone removed from the sampling sites.

#### Next generation sequencing analysis

Total DNA extracts from bone were sent to Hudson Alpha Institute of Biotechnology Genome Services Laboratory (Huntsville, AL) for sequencing of the V3-V4 region of the 16S rRNA gene and V4-V5 of the 18S rRNA gene using 300 PE chemistry on an Illumina MiSeq instrument. Library preparation was performed by Hudson Alpha according to Illumina protocols. Primers included S-D-Bact-0564-a-S-15 and S-D-Bact-0785-b-A-18 for the 16S rRNA gene [41] and 574f and 1132r for the 18S rRNA gene [42]. Raw sequence data is available at NCBI Sequence Reach Archive, Accessions PRJNA540930 and TBD.

Adapters were removed by Hudson Alpha prior to data distribution. Read quality was assessed using fastqc (v. 0.11.7) and multiqc (v. 1.5). Primers were removed using cutadapt (v. 1.14) [43], and reads were quality trimmed using trimmomatic (parameters: LEADING:15 TRAILING:10 SLIDINGWINDOW:4:20 MINLEN:15) (v. 0.36) [44]. Data were further trimmed, aligned, and classified using mothur (v. 1.39.5) according to the mothur SOP [45]. 16S rRNA and 18S rRNA sequences were aligned and classified into operational taxonomic units (OTUs) at 97% sequence identity, using SILVA (v. 128). Statistical analyses and visualizations were conducted in R (v. 3.4.1) [46], primarily using phyloseq (v.1.20.0) [47] and dependencies. Mothur code, R code, and associated files, including metadata, can be found at https://github.com/aemmons90/Surface-Bone-Microbe-Project.

Samples with less than 5,000 reads were removed from analyses, and remaining samples were rarefied to even depth by the smallest library (16S rRNA min. library = 48,288 reads; 18S rRNA min. library = 5,368 reads) prior to alpha and beta diversity measurements including ordination methods and visualizations based on ordination methods (S1 and S2 Figs). Bray-Curtis dissimilarities were computed for all ordinations. Alpha diversity metrics including Inverse Simpson and observed richness were computed using a subsampling approach, in which richness and diversity metrics were computed for a total of 100 iterations, each scaled to even depth.

#### Sequence quality analysis

Two samples failed to sequence using 16S rRNA primers, while twenty samples failed to sequence using 18S rRNA primers. Fastqc and multiqc demonstrated high quality reads in the forward direction, with a drop in mean quality Phred scores in the reverse direction at an approximate base pair position of 200 (Phred Score < 25). Following cutadapt and trimmomatic, total 16S rRNA contigs were reduced by 46%. This was further reduced by an additional 14% following further processing in mothur, resulting in a total read loss of 60% (from 37,185,525 to 14,958,201 sequences). This left a total of 14,958,201 sequences, of which 692,709 were unique.

18S rRNA sequences presented an additional challenge; using 300 PE chemistry, forward and reverse reads overlapped by ~59 base pairs (bp) (See [48]). Fastqc and multiqc showed a significant reduction in mean base quality in both forward and reverse reads. Forward reads showed a drop in mean quality scores at an approximate position of 250 bp (Phred scores < 25), the same drop in quality was observed in reverse reads at ~200 bp. As a consequence, trimming to remove low quality base pairs resulted in a dramatic loss of reads. Following cutadapt and trimmomatic, total 18S rRNA contigs were reduced by 46%, and after further processing in mothur, sequences were further reduced by 49%, resulting in a total read loss of 95% (from 30,253,173 sequences to 1,518,971). This left a total of 1,518,971 sequences, of which 181,486 were unique. Due to poor read quality, individual A was removed from additional data analysis in phyloseq, resulting in a remaining 7,901 OTUs across 91 samples. Following the removal of samples with less than 5,000 reads, a total of 71 samples remained.

#### Data analysis

All data analyses, excluding random forests tests, were conducted in R (v.3.4.1). Two-factor analysis of variance tests (ANOVAs) were used to examine differences in log transformed human DNA concentrations by individual and body region (i.e., head, upper torso, arm, hand, lower torso, leg, foot). Assumptions such as normality and homogeneity of variance were tested using D’Agostino’s normality test (package = fBasics v. 3042.89) [49] and Levene’s test (package = car v. 3.0.2) [50], respectively. Regression analysis was then used to assess the relationship between human DNA concentrations from bone samples and hypothesized predictor variables (i.e., bacterial DNA gene abundances, total DNA concentrations, and percentage cortical content). Human DNA concentrations, bacterial gene abundances, and total DNA concentrations were log-transformed prior to linear regression. Multiple regression analysis was also performed, treating log transformed human DNA as the dependent variable and log transformed bacterial gene abundances, log transformed total DNA, and percentage cortical content as independent variables, including their various interactions. Assumptions including heteroskedasticity, normality, autocorrelation, and multicollinearity were tested using the R package sjstats (v. 0.17.0) [51].

Kruskal-Wallis tests were used to assess statistical significance in alpha diversity metrics, followed by multiple comparisons with false discovery rate (FDR) adjusted p-values. Permutational multivariate analysis of variance tests (PERMANOVAs), applying 999 permutations, were used to assess statistical significance in beta diversity between categorical variables of interest including body region, individual (A, B, and C), human DNA category, and cortical category. These same variables were tested for homogeneity of multivariate dispersion, using 999 permutations. Human DNA category was an arbitrary categorical variable created by dividing a continuous variable, human DNA concentration, by quartiles in each dataset, each quartile defining a category used for factor analysis. Cortical category was established by using the mean percentiles of cortical bone composition at each sampling site as described above [0 (teeth), 1 (80 to 100%), 2 (70 to 79%), 3 (50 to 59%), 4 (40 to 49%), 5 (30 to 39%), 6 (<39%)]. However, because no bones comprised the 60-69% category, this category was eliminated for the purpose of data analysis. In addition, the frontal bone was assigned to the third category rather than the fourth, due to the mean being affected by a single individual. SIMPER, similarity percentages, followed by non-parametric Kruskal-Wallis tests with FDR corrected p-values, were used to determine OTUs significantly contributing to differences between individuals and human DNA category (seq-scripts release v. 1.0) [52]. Random forest models were generated using Python (v. 3.5.2) and scikit-learn (v. 0.19.2) [53] to identify OTUs contributing to human DNA preservation patterns. OTUs were merged at the genus level, and all samples were used to generate the model (bacteria, n = 162; microbial eukaryotes, n = 71; combined datasets, n = 71). Data were randomly split into training (3/4) and testing (1/4) sets.

## Results

### Bacterial and human quantification via qPCR

Though bacterial gene abundances, which were used as a proxy for bacterial loading, were often high when human DNA quantities were low, for example in the teeth, upper torso, lower torso, and in the hand, this relationship was not consistent across all body regions. Despite foot bones having some of the highest human DNA quantities, these also corresponded with high bacterial gene abundances (Fig 1). While bones with high cortical content generally demonstrated lower bacterial infiltration, bacterial gene abundance was not a significant predictor of percent cortical content (adjusted R^2^ = −0.03) (Fig 2A). Total DNA was, however, a significant predictor of percent cortical content *(p* < 0.001, F = 71.43, DF = 1, 33, adjusted R^2^ = 0.67); as the percentage of cortical bone decreased, total DNA increased (Fig 2A).

**Fig 1.**
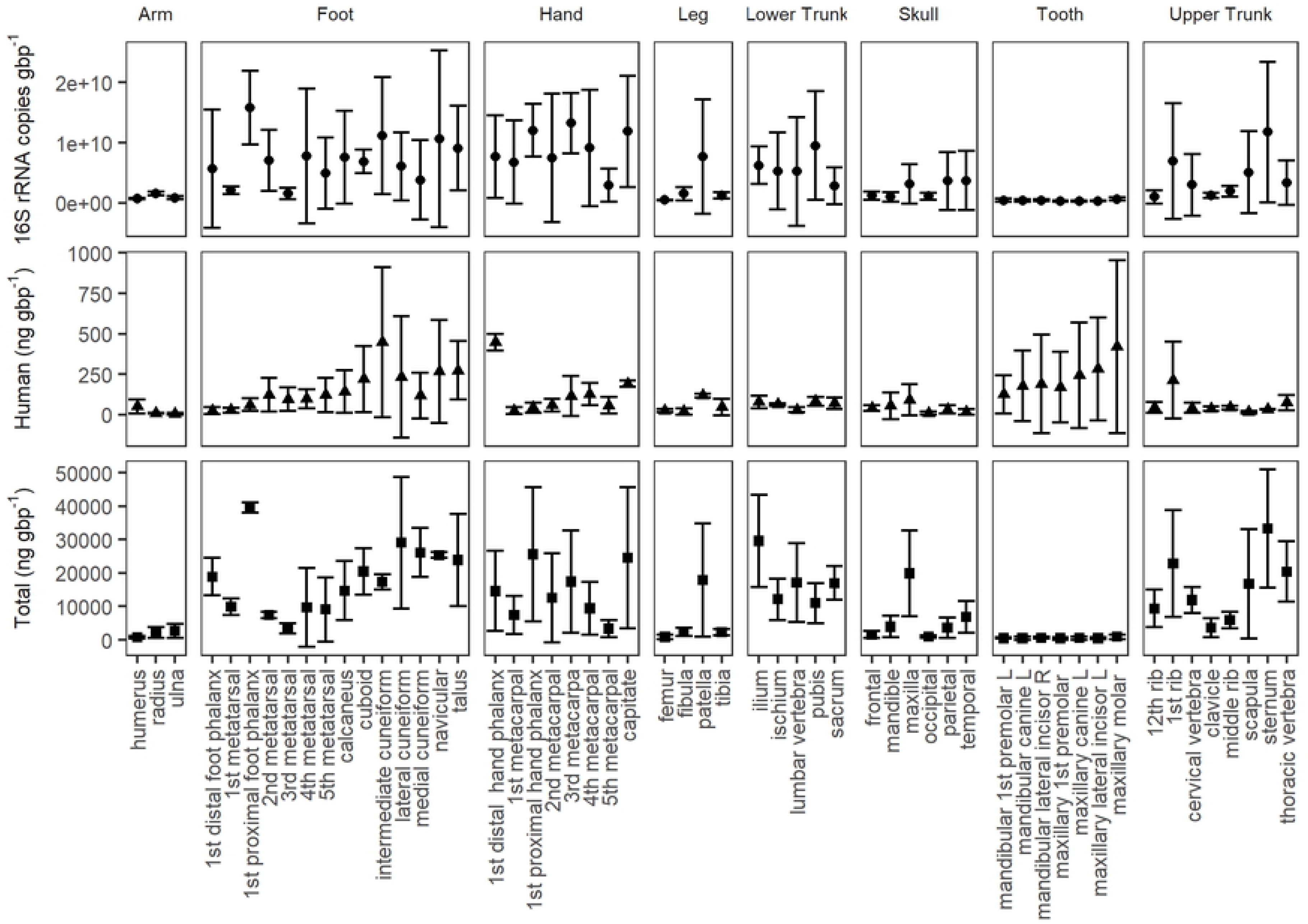
Mean total DNA (Total (ng gbp^-1^)) or the concentration of DNA extracted, mean human DNA concentration (Human (ng gbp^-1^), as quantified using qPCR, and bacterial gene copies (16S rRNA copies gbp^-1^), quantified using qPCR by bone type (n = 3 individuals). Concentrations are presented as nanograms (ng) per gram of bone powder (gbp^-1^). Bars represent standard deviations where n = 3.

**Fig 2.**
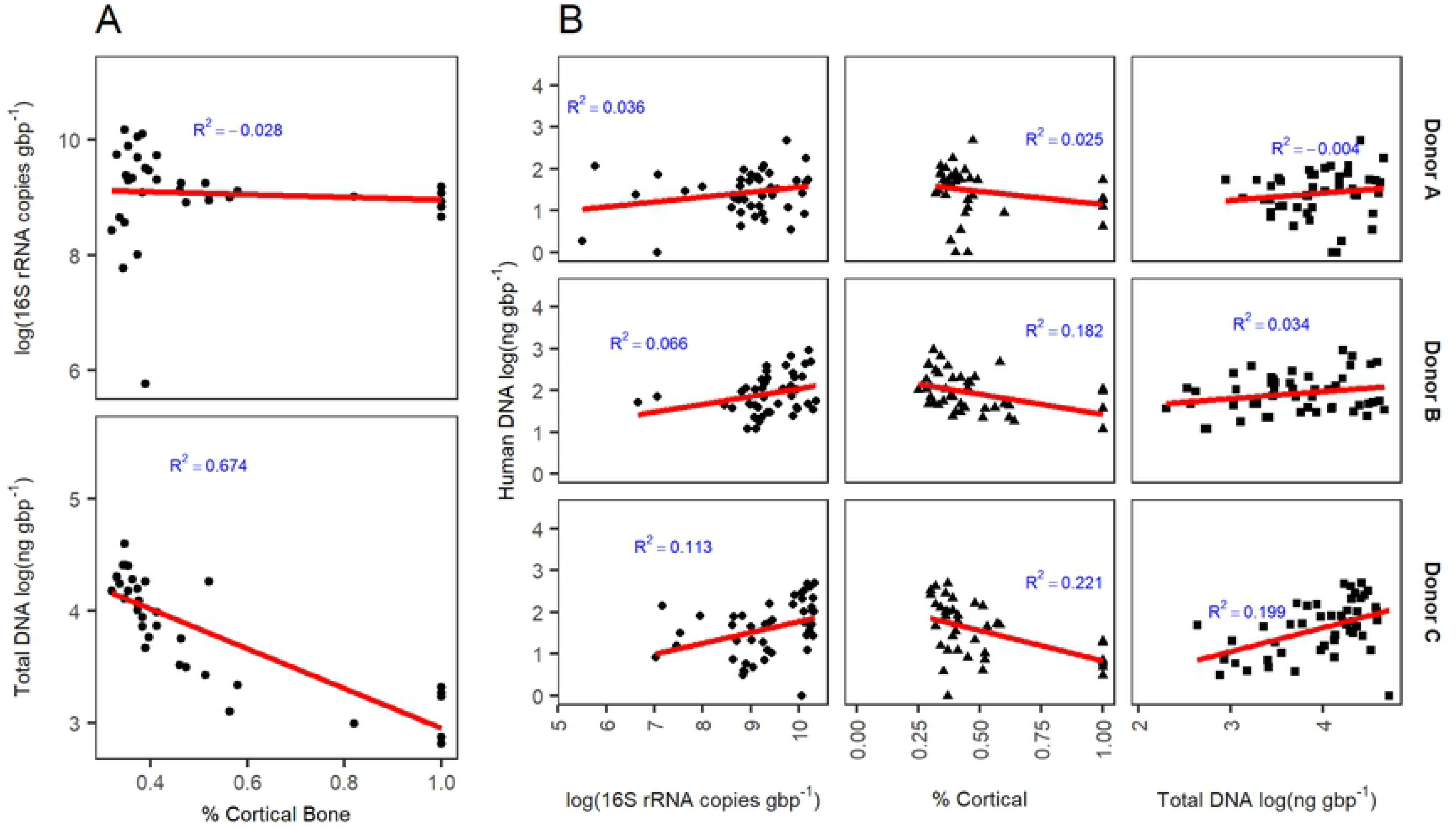
(A) Percent cortical content compared with log-normalized bacterial gene abundances and log-normalized total DNA, averaged by bone type (n = 3). (B) Bacterial gene abundances, percent cortical content, and total DNA compared with human DNA concentrations by individual (A, B, C). Raw data is shown. The red line demonstrates the best fit linear regression.

When excluding teeth, human DNA quantities were significantly different by individual *(p* < 0.001, F = 12.06, DF = 2) and body region (DF = 6, *p* < 0.01, F = 4.52), with a significant interaction between body region and individual (*p* < 0.05, F = 1.87, DF = 12). On average, individual B had greater concentrations of human DNA than C or A, with individual A having the lowest concentrations. Therefore, to test the effects of various predictor variables on human DNA recovered from bone, individuals were assessed independently. Bacterial gene abundance did not significantly predict human DNA concentration in two out of three individuals (A, *p* = 0.12; B, *p* = 0.05), while bacterial gene abundance demonstrated a positive relationship with human DNA concentration in individual C (*p* = 0.01, F = 6.85, adjusted R^2^ = 0.113) (Fig 2B). A similar relationship was observed for total DNA, which showed a positive relationship with human DNA concentration for individual C *(p* = 0.001, F = 12.41, adjusted R^2^ = 0.199). In addition, percent cortical content was a significant predictor of human DNA concentration in two of three individuals (B, *p* = 0.003, F = 10.33, DF = 1, 41, adjusted R^2^ = 0.182; C, *p* = 0.002, F = 11.75, DF = 1, 37, adjusted R^2^ = 0.221).

When including all predictors (i.e., bacterial gene abundance, total DNA concentration, and percent cortical content) in a single model, the assumption of multicollinearity was not met, indicating that predictor variables were highly correlated.

### Bacterial community analysis

Bacterial communities showed contributions from 47 phyla; of these, only 12 demonstrated greater than 2% relative abundance when averaged by bone type: Proteobacteria (20 to 57%), Actinobacteria (4 to 37%), Firmicutes (2 to 35%), Bacteroidetes (2 to 21%), Planctomycetes (0.2 to 11%), Saccharibacteria (0.2 to 12%), Chloroflexi (2.8 to 7.8%), Verrucomicrobia (0.05 to 4.7%), Chlamydiae (0.02 to 3.9%), Acidobacteria (0.04 to 2.2%), BRC-1 (0.009 to 2.3%), and Deinococcus-Thermus (0 to 7.0%) (Fig 3).

**Fig 3.**
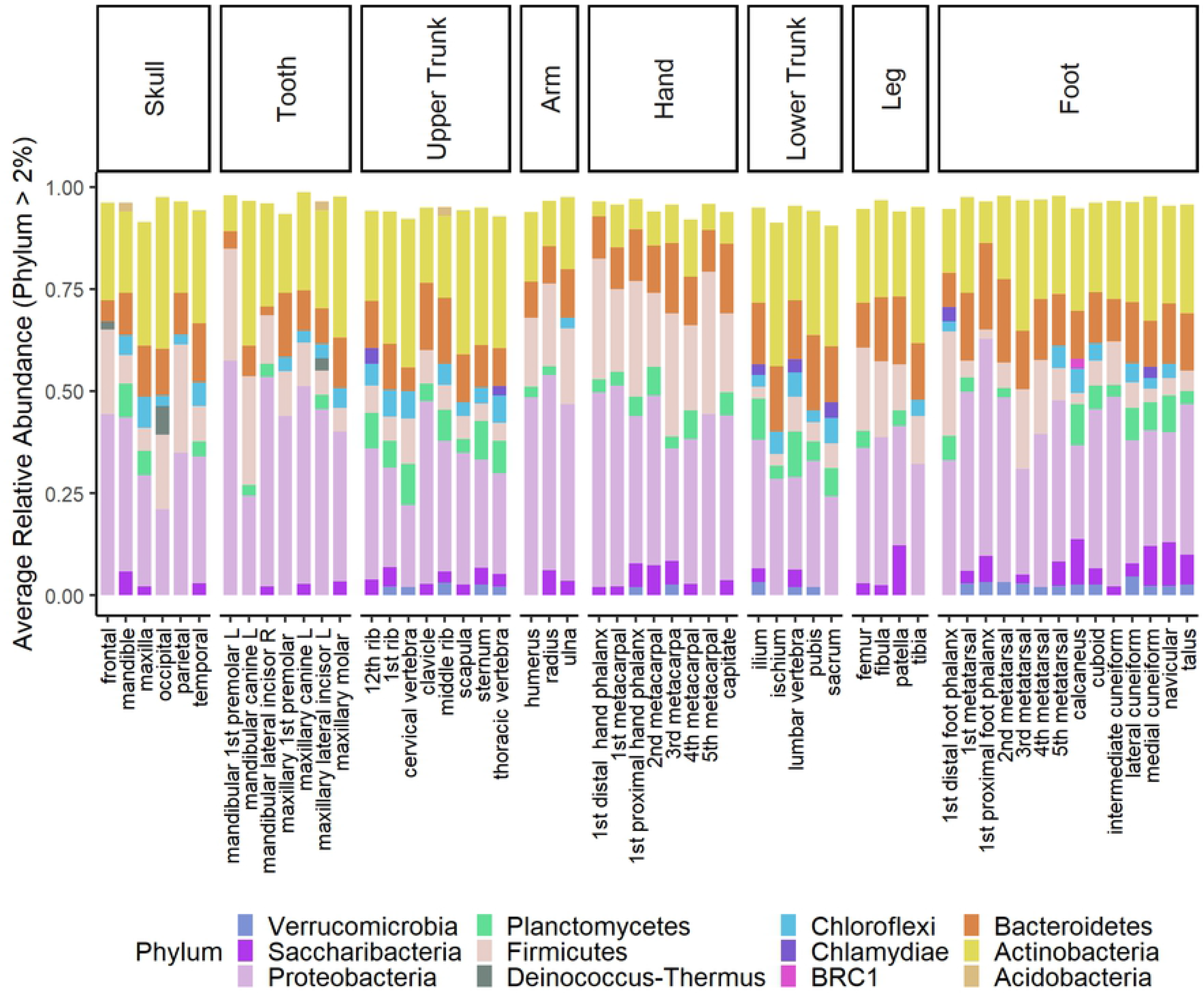
Bacterial phylum-level community membership. Mean relative abundances greater than 2% for all individuals combined. Bone phyla membership was averaged by bone type (n = 3), except in the navicular, occipital, and sternum (n = 2).

Bacterial communities significantly differed by individual (*p* = 0.001, F = 11.08, DF = 2) (Fig 4), body region (*p* = 0.001, F = 3.99, DF = 7), human DNA concentration (*p* = 0.02, DF = 3, F = 1.48), and cortical bone content (*p* = 0.003, F = 1.28, DF = 5). There was a significant interaction between body region and individual (*p* = 0.001, F = 2.70, DF = 14) and body region and cortical content (*p* = 0.02, F = 1. 23, DF = 4) (Fig 5). Heterogeneous multivariate dispersion was observed by individual (*p* = 0.016), body region (*p* = 0.001), and cortical category (*p* =0.001), but not human DNA (*p* = 0.27); bacterial communities from individual A and C clustered more tightly compared with individual B (Fig 4). When examining individuals independently, body region remained significant (A, *p* = 0.001; B, *p* = 0.001; C, *p* = 0.001), while cortical content remained significant in individuals B and C (B, *p* = 0.003; C, *p* = 0.03) but not A (*p* = 0.63).

**Fig 4.**
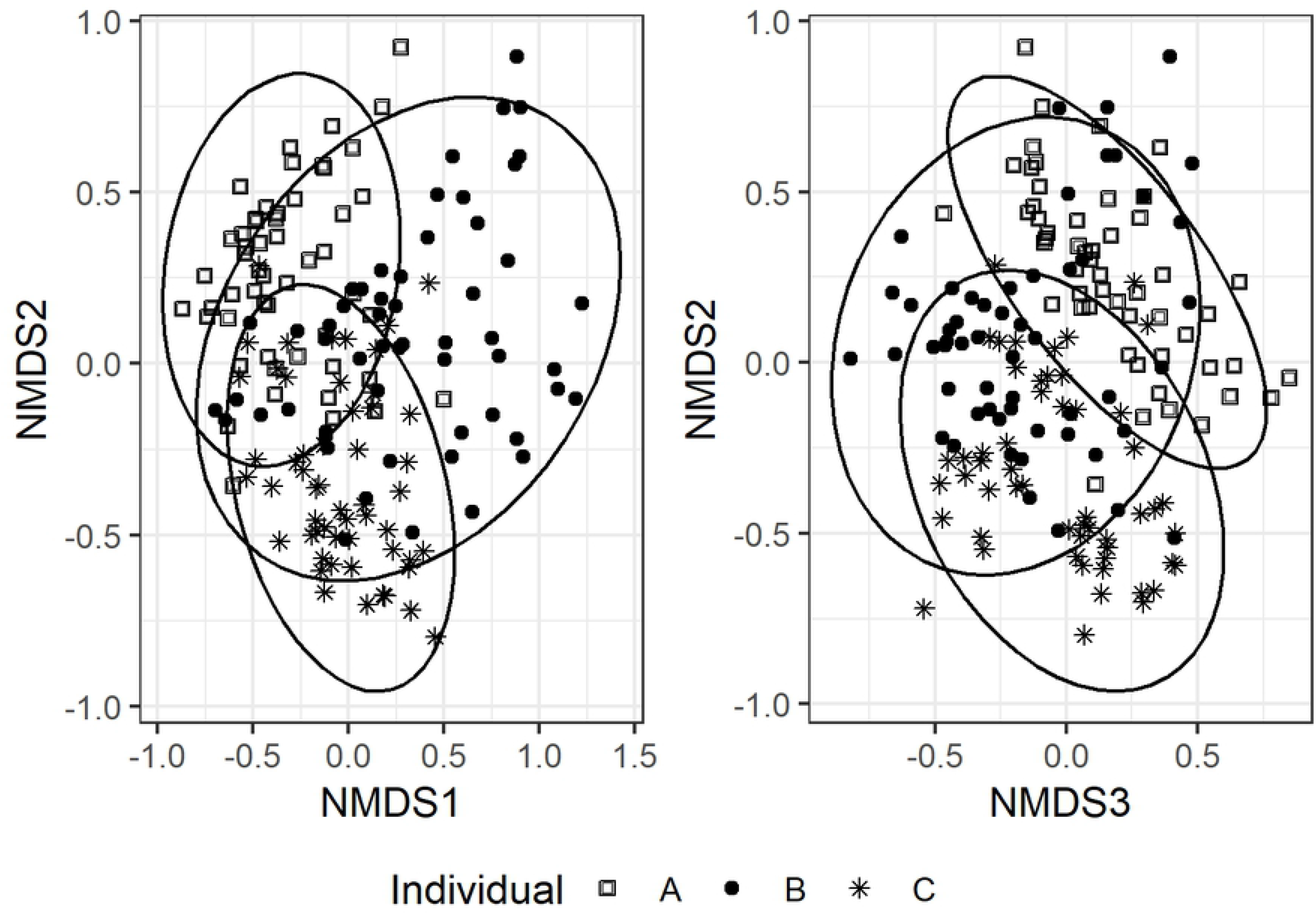
Non-metric multidimensional scaling (NMDS) ordination performed on Bray-Curtis dissimilarities of bone bacterial communities (n =162) and visualized by individual. Stress = 0.14 and k = 3; ellipses represent 95% confidence intervals.

**Fig 5.**
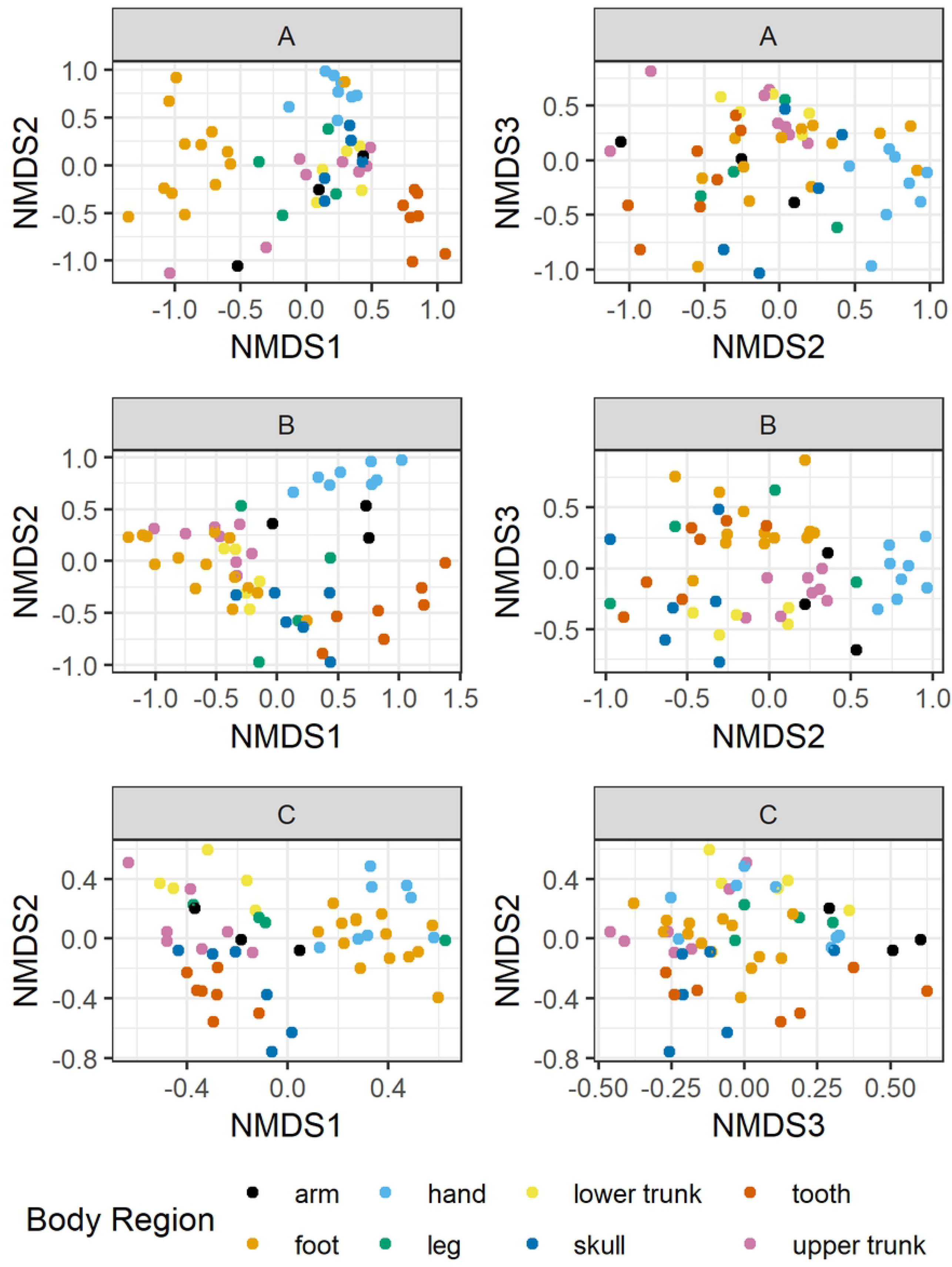
Non-metric multidimensional scaling (NMDS) ordinations on Bray-Curtis dissimilarities of bone bacterial communities. Ordinations were conducted independently by individual (A: n = 53, B: n = 55, C: n = 54) and visualized by body region (A: stress = 0.14, k = 3; B: stress = 0.10, k = 2; C: stress = 0.10, k = 3). The letters “A”, “B”, and “C” above figure panels refer to individuals.

Diversity was significantly different by individual (*p* < 0.01, DF = 2); individual A had the lowest diversity (mean = 30.0), while individual C had the greatest diversity (mean = 46.9) (S3 Fig). When each individual was considered independently, diversity also significantly differed by body region (A: *p* < 0. 01, X^2^ = 19.0, DF = 7; B: *p* < 0.0001, X^2^ = 34.6, DF = 7); C: *p* < 0.001, X^2^ = 24.9, DF = 7) (S3 Table; S4 Fig). Body regions from A followed a different trend in diversity than B or C. Richness did not show significant differences by individual (*p* > 0.05, X^2^ = 3.97, DF = 2), but did significantly differ by body region (*p* < 0.0001, X^2^ = 46.0, DF = 7). Observed richness was greatest in the upper and lower torsos (S5 Fig).

OTUs driving differences between individuals included predominantly soil taxa from the following families: Streptosporangiaceae, Nocardiaceae, Comamonadaceae, Pseudomonadaceae, Xanthomonadaceae, Clostridiaceae, Brevibacteriaceae, Streptomycetaceae, Intrasporangiaceae, unclassified Thermomicrobia, and Mycobacteriaceae. Notably, OTUs identified as *Simplicispira* and an unclassified member of Streptosporangiaceae were found at greater abundances in A, while *Stenotrophomonas* and *Rhodococcus* showed greater abundances in individual C. *Brevibacterium,* an unclassified member of Thermomicrobia, and *Pseudomonas* were greatest in B (S6 Fig). Although three OTUs significantly contributed to differences by human DNA category (two *Streptomyces* and one *Mycobacterium),* these OTUs did not remain significant after correcting p-values using FDR.

Random forest models were used to identify bacterial OTUs associated with differences in human DNA concentrations. The initial model generated had a mean absolute error of 91.7 (*p* = 0.03, adjusted R^2^ = 0.09), with 30 predictor OTUs identified (S7 Fig). Important predictor OTUs were represented by Actinobacteria (importance = 30%), Bacteroidetes (17%), Firmicutes (23%), and Proteobacteria (30%). Contributing OTUs greater than 1% included the genera *Clostridium,* unclassified Dermacoccaceae, *Paracoccus*, and *Actinotalea* (S7A Fig). The model only slightly improved when excluding teeth from the analysis (mean absolute error = 72.5, *p* = 0.02, adjusted R^2^ = 0.12). When teeth were excluded, the top five predictor OTUs became unclassified Dermacoccaceae (14%), unclassified Desulfuromonadales (9%), *Clostridium* (3%), unclassified Gaiellales (3%), and unclassified Mollicutes (3%).

### Microbial eukaryotic community analysis

Microbial eukaryotic communities showed large contributions from Ascomycota (mean relative abundance 40%), Apicomplexa (21%), Annelida (19%), Basidiomycota (17%), Ciliophora (14%), and enigmatic Eukaryota (including *Incertae sedis*) (14%), with additional contributions from Cercozoa (9%) Peronosporomycetes (8%), Nematoda (7%), and Cryptomycota (6%). Unclassified Eukaryota had a mean contribution of 8% (Fig 6). While Apicomplexa had a high mean relative abundance (21%), this was dominant in a single sample, a fibula from individual B.

**Fig 6.**
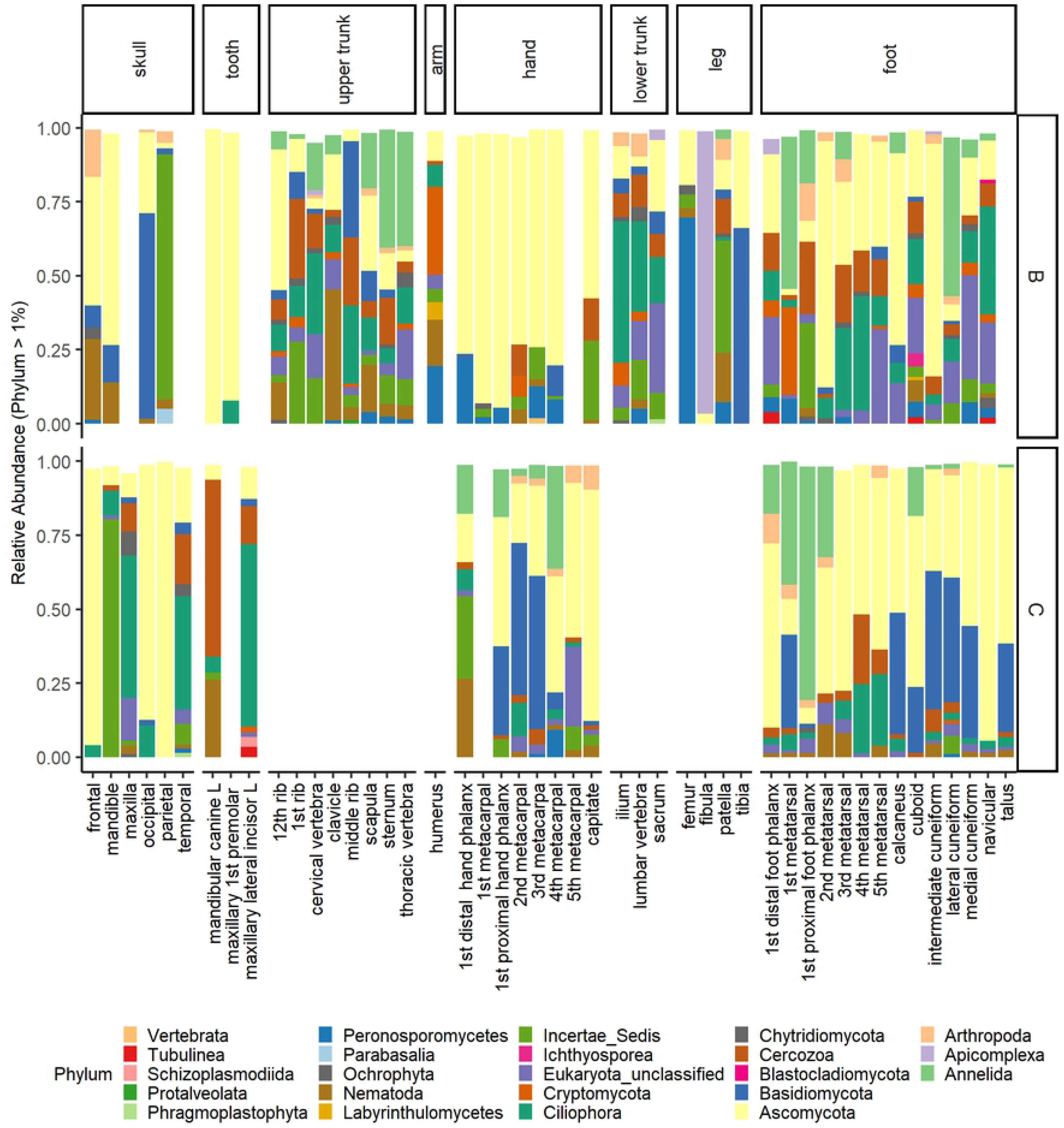
Relative abundance of eukaryotic phyla by bone type and individual. Relative abundance is shown for only those phyla with greater than 1% relative abundance, and for two of the three individuals (B and C). Data were not averaged by bone type.

Eukaryotic communities showed similar patterns in beta diversity compared to bacterial communities. When testing differences between body region, individuals, human DNA quartiles, and cortical content, microbial eukaryotic communities significantly differed by individual (*p* = 0.001, F = 8.69, DF = 1), body region (*p* = 0.001, F = 2.83, DF = 7), human DNA (*p* = 0.02, F = 1.41, DF = 3), and cortical content (*p* = 0.001, F = 1.60, DF = 5), with a significant interaction between body region and individual (*p* = 0.001, F = 4.08, DF = 3), body region and human DNA (*p* = 0.02, F = 1.25, DF = 4), and individual and cortical content (*p* = 0.008, F = 1.72, DF = 1). Due to sequence loss, alpha diversity metrics were not computed.

A random forest model was also applied to the microbial eukaryotic dataset to identify OTUs contributing to patterns of human DNA preservation. The resulting model was not significant, with a mean absolute error of 171.96 (*p* = 0.14, adjusted R^2^ = 0.07). The most important predictor taxon identified (OTU0003), contributing to 33% of the model, was an unclassified Saccharomycetales. Eukaryotic and bacterial OTU data were combined and a random forest model was constructed for shared samples to predict human DNA concentrations. The resulting model was significant (mean absolute error = 175.71, *p* = 0.03, adjusted R^2^ = 0.21). Again, the top predictor taxon was OTU0003 with an importance value of 10% or 0.1; this Saccharomycetales OTU decreased in abundance in the skull of individual B as human DNA concentrations increased (S8 Fig). Other important contributors, with importance values greater than 1% or 0.01, included bacterial genera from the phyla Actinobacteria (*Microbacterium,* 6%, Gaiellales uncultured, 5%, *Leifsonia,* 2%, *Williamsia,* 2%), Proteobacteria (*Stenotrophomonas*, 2%), Firmicutes (Clostridiales Family XI uncultured, 2%), Gemmatimonadetes (unclassified Gemmatimonadaceae, 2%), and Planctomycetes (*Zavarzinella,* 2%).

## Discussion

### Characterizing the postmortem bone microbiome

The post-mortem bone microbiome is diverse and variable in the human skeleton two years after death. Excluding Planctomycetes and Saccharibacteria, dominant taxa observed in this study were also shown to dominate human rib samples from twelve individuals that had decomposed at the ARF [37]. Rib samples from the current study most closely resembled dry remains from Damann et al. [37] in phyla-level contributions, but also contained taxa proportions greater than 2% from Verrucomicrobia, Saccharibacteria, Planctomycetes, Chloroflexi, and Chlamydiae (S10 Fig). Discrepancies in observed taxa may be due to differences in sample size and sequencing analysis methodologies. Planctomycetes, a phylum commonly associated with aquatic environments, and Saccharibacteria, a phylum containing multiple environmental taxa have been observed in gravesoils [54,55].

Ascomycota, observed in 100% of samples (71 of 71) in the eukaryotic dataset, and Basidiomycota, observed in 55% of samples, were the dominant microbial eukaryotes. This was unsurprising, as these fungal phyla contain multiple saprophytic groups that have previously been observed in association with decomposing carrion [56]. In addtion to fungi, multiple phyla of protists were also detected, including Apicomplexa (5 of 71 samples), Ciliophora (49 of 71 samples), and Cercozoa (49 of 71 samples). Protists found in association with bones may be opportunistic, potentially transferred to remains via soil, scavengers, insects, precipitation, and run-off, and may be active fungal and bacterial consumers. For example, the genus *Rhogostoma,* which was prevalent in samples from individuals A and B, is known to consume both fungal and bacterial species [57]. Similarly, Nematoda were detected, with the majority of sequences belonging to the family Rhabditidae, which contains bacterivorous members, previously observed in decomposition research [58–61]. Other bacterivores detected within human bones included Tubulinea, Cercozoa, and Apicomplexa, which have also been found in soils underlying human remains [61]. Cercozoa and other testate amoeba are extremely sensitive to environmental change, and generally decrease in soil with cadaveric inputs [62]. While certain species have responded with positive growth during late stage decomposition (from 1 month to 1 year postmortem) [63], their presence in bones over a year after death likely reflects a shift back to more oligotrophic conditions.

Presence of Deinococcus-Thermus, a phylum well-represented by thermophiles [64], at greater than 1% relative abundance in 6% of samples, is suggestive of a harsh environment. Bones deposited on the soil surface are exposed to daily and annual temperature contrasts. East Tennessee experiences freezing winter temperatures and temperatures greater than 37°C in the summer, which can influence moisture availability. As indicated by Reeb et al. [38], bone may provide shelter from harsh environments (i.e., variable temperature, UV). Individual C had greater abundances of Deinococcus-Thermus than B and A (S11 Fig), likely due to the greater duration of exposure to environmental fluctuations, including temperature and precipitation (S9 Fig). The majority of samples with abundances greater than 1% were from the skull including cranial elements and teeth. The cranium is often one of the first anatomical regions to skeletonize during decomposition due to low tissue biomass and high larval presence [65] and likely experiences greater intervals of environmental exposure.

### Community differences by individual and anatomical region

Beta diversity analyses showed differences in bone microbial communities, including both prokaryotes and eukaryotes, by individual and body region. This is unsurprising, as there is extensive research on the living human microbiome and the multitude of variables leading to differences in microbial community structure and composition between individuals including life history (e.g., health and diet) [66–68]. Two of the three individuals had a history of diabetes (individuals A and C), which may have contributed to differences in microbial community structure and composition [69]. Moreover, placement duration at the ARF and differences in temperature and precipitation likely contributed to differences observed between individuals. In particular, bacterial alpha diversity was lowest in individual A and greatest in C, reflecting differences in exposure duration (S3 and S9 Figs). The impact of soil microbiota is expected to increase overtime with prolonged soil contact [37]. Because of this, we hypothesize that differential rates in skeletonization likely influence bone microbial composition and structure at any given time point, which likely has implications for post-mortem interval estimation.

Recently, Pechal et al. [70] showed microbial differentiation by anatomic region (i.e., external sites from the auditory canal, eyes, nose, mouth, umbilicus, and rectum) up to 48 hours after death. Though they speculated that this pattern would likely attenuate with longer post-mortem intervals, this has yet to be tested. Here, bone microbial communities retained differences by anatomic location in individuals with post-mortem intervals greater than 1 year. Micro-environmental differences in soil communities as well as differences in enteric microorganisms and their abilities to compete and persist with soil microorganisms colonizing the body likely contributed to spatial differences observed in anatomic regions and between different individuals. Research on the human microbiome has shown microbial community uniqeness by individual as well as body site and time [71,72], and has recently gained utility in forensics [73,74].

Nicholson et al. [75] demonstrated that bones in similar environments showed drastic differences in bone preservation, despite similarities in soil pH and drainage. This evokes the question: if not the environment, then what is the source of these differences? Enteric/putrefactive bacteria have been posited as the primary source of microbial bone degradation in pig remains; neonatal pig remains demonstrated no evidence of microbial degradation, which researchers hypothesized as being related to the relative sterilitiy of infant guts compared with adults [31]. While the source of bacteria in this study remains unknown, as we have no gut or soil samples prior to placement to track bacterial translocation, we suspect that both soil and gut microbes are able to colonize and aid in bone degradation (e.g.,[76]). We have previously demonstrated that human-associated *Bacteroides,* an obligate anaerobic member of the human gut microbiome, can persist for long time periods in soils impacted by decomposing human remains [54,59], providing evidence that these gut microbes are transferred to the environment and have the potential to colonize bone. The extent to which enteric microorganims are able to move throughout the body post-mortem is likely limited, and distance from the gut may be a crucial factor controlling differences in microbial communities by body region. However, Pechal et al. [70] recently observed an increase in gene abundance associated with bacterial motility during decomposition, so this area of postmortem microbiology merits further study.

Bone microstructure (i.e., the percentage of cortical content) also influenced differences in microbial communities. Communities differed by cortical bone percentage likely due to the presence of greater void space in cancellous bone compared with cortical bone, facilitating ease of invasion, especially for incidental taxa or soil contaminants (e.g., potentially Verrucumicrobia). However, this may also be related to nutritive differences; cancellous elements may harbor more labile remnant material such as red marrow [22], while cortical bone may be considered more recalcitrant. This may account for patterns observed in total DNA concentrations and bacterial gene abundances. Bacterial gene abundance was not a significant predictor of human DNA concentration, and cases where bacterial gene abundance did significantly predict human DNA (i.e., individual C), the relationship was positive, indicating that the degree of microbial loading does not negatively impact the pattern of skeletal DNA preservation in remains with environmental exposure up to two years. Rather, the presence of specific taxa likely has a greater impact on skeletal integrity.

Additionally, presence of both aerobic and anaerobic genera points to the existence of micro-spatial differences within a single bone. This phenomenon is also observed in soils where anaerobic microsites can persist within a well-drained, well-aerated soil. Extracellular polymeric substances were observed surrounding living cells on bison bone at Yellowstone National Park [38]. This highlights the importance of biofilm production in microbial bone colonization. Though microscopy was not performed here to confirm biofilm presence, we hypothesize that biofilm production combined with increased microbial biomass during decomposition plays an important role in the development of micro-spatial differences in oxygen access and respiration strategies.

### Microbial taxa associated with skeletal DNA preservation

Random forest models were minimally successful in identifying microbes related to DNA preservation patterns, however models were likely complicated by microbial community differences by individual and body region. While bacterial OTUs produced more accurate random forest models than eukaryotic OTUs, the best model resulted when combining both bacterial and eukaryotic data sets, with a Saccharomycetales OTU identified as the most important contributor to the model. Saccharomycetales, commonly associated with the oral microbiome of healthy humans [77], decreased in abundance with increased human DNA concentrations in the cranium of individual B. Oral microbes may persist throughout decomposition and may be implicated in DNA survival.

Similarly, bacterial random forest models were conflated by body region; genera *Actinotalea* and *Paracoccus*, showed increased abundances with human DNA concentrations in teeth, while Dermacoccaceae demonstrated increased abundances in feet. Importantly, increased abundances of *Clostridium,* a genus that contains known collagenase producers [12], were associated with decreased human skeletal DNA concentrations. The foot is the farthest anatomical region from the gut, and interestingly, bones of the feet had some of the highest human DNA concentrations. If the *Clostridium* present in bones is primarily derived from the gut, then distance from the gut may be an important factor related to human DNA degradation. Though predictor taxa could be identified using random forest models, their functional role in DNA degradation, if any, remains unclear. The variation seen by body region and individual may be minimized by increasing the research sample size to include more individuals.

## Conclusions

Most of what is known regarding the microbial degradation of bone is from histological research concerning archaeological bone (e.g., [32,34,78–80]). The current study used next generation sequencing technologies to provide a survey of bacteria, fungi, and protists potentially capable of bone colonization. Though specific taxa were correlated to patterns of human DNA preservation using random forest models, the functional role of identified bone microbes remains unknown. Because the target of this study was DNA, which provides information regarding presence rather than activity, it is difficult to discern incidental taxa, i.e. taxa that are present and inactive, from taxa that are actively degrading bone. This is a longstanding challenge in microbial ecology: linking structure and function. Remnant extracellular DNA of microbial origin is a problem [78], and microbial DNA can bind to hydroxyapatite similar to human DNA [79, 80], further complicating observed differences in community composition and structure. Nevertheless, the current study presents a first step in characterizing microbial community differences across bone types within and between individuals following skeletonization. Ultimately, this provides a foundation for understanding the postmortem colonization of bone by microbes and the subsequent effects on bone stability and human DNA preservation and may help guide targeted human DNA recovery.

## Acknowledgements

Thanks to Mundorff and DeBruyn lab members for their support and guidance in this research. In addition, the authors would like to thank Dr. Yong Bradley for providing access to the clinical CT scanner at the University of Tennessee Medical Center, and to Shelley Acuff for operating the CT system and assisting with bone set-up prior to scanning.

## Supporting Information

**S1 Table. Donor Information for the three individuals placed at ARF 2009.**

**S2 Table. Bones sampled from each of the three individuals (Mundorff and Davoren 2014).**

**S1 Fig. 16S rRNA targeted metagenomics read distribution**. The minimum library size was 48,288 reads, while the mean library size was 92,334.6 reads; the maximum library was 150,228 reads.

**S2 Fig. 18S rRNA targeted metagenomics read distribution**. The minimum library size was 5,368 reads; the mean library size was 19,853 reads, and the maximum library size was 40,977 reads.

**S3 Fig. Bacterial alpha diversity (Inverse Simpson) and richness (observed) by individual.** **Indicate significance below an alpha value of 0.01. No significance is denoted by “ns”.

**S3 Table. Bacterial alpha diversity metrics including observed richness and diversity (Inverse Simpson)**. Data was separated by individual, and the mean and standard deviation was computed by body region. Significance levels are represented by asterisks (*p < 0.05; **p < 0.01); multiple comparison tests by individual were only conducted using Inverse Simpson indices. Lower case letters in parenthesis refer to body regions. A body region with an exponent corresponding to another body region indicates that those two regions have significantly different diversity indices.

**S4 Fig. Inverse Simpson (diversity) calculated from the bacterial dataset**. Individuals (“A”, “B”, and “C”) were assessed independently

**S5 Fig. Richness (observed) calculated from the bacterial dataset; individuals were combined**

**S6 Fig. SIMPER results by individual (A: n=53, B: n=55, C: n=54)**. Results include only those OTUs that demonstrated a significant difference by individual following SIMPER. All OTUs of a given genus are not represented here. The x-axis contains information on the taxonomic identification of these OTUs at the family level.

**S7 Fig. Random forest regression**. (A) Bacterial OTUs important for predicting human DNA concentrations. Importance, as a value between zero to one, is represented in color. Human DNA concentrations greater than 400 ng gbp^-1^ are labeled by body region. Abundance refers to relative abundance by bone sample, represented by values zero to one. (B) Model accuracy as a function of test values versus predicted values. The solid red line represents the modeled data; the dotted line represents an expected model if 100% accuracy.

**S8 Fig. Relative abundance of the unclassified Saccharomycetales OTU plotted against human DNA concentration.** The label “B” refers to individual B.

**S9 Fig. Changes in temperature and precipitation for the duration of deposition of each donor.** Donors are labeled “A”, “B”, “C”, and “ADD” refers to accumulated degree days, an indicator of both time and temperature. Data obtained from NOAA (https://www.noaa.gov/).

**S10 Fig. Phylum-level bacterial community membership in human rib samples**. Relative abundance was averaged by bone type, combining results from three individuals (n = 3). Only taxa with average relative abundances greater than 1% are shown.

**S11 Fig. Bacterial community phylum-level contributions visualized by individual (“A”, “B”, and “C”) and body region (arm, foot, hand, leg, lower trunk, skull, tooth, upper trunk)**. Only relative abundances greater than 1% are displayed.

## References

1. Megyesi MS, Nawrocki SP, Haskell NH. Using accumulated degree-days to estimate the postmortem interval from decomposed human remains. J Forensic Sci. 2005;50: 618–626. doi:10.1520/JFS2004017

2. Keenan S. From bone to fossil: a review of the diagenesis of bioapatite. Am Mineral. 2016; doi:10.2138/am-2016-5737

3. Perry WL, Bass WM, Riggsby WS, Sirotkin K. The autodegradation of deoxyribonucleic acid (DNA) in human rib bone and its relationship to the time interval since death. J Forensic Sci. 1988;33: 144–53. Available: http://www.ncbi.nlm.nih.gov/pubmed/3351452

4. Elsner J, Schibler J, Hofreiter M, Schlumbaum A. Burial condition is the most important factor for mtDNA PCR amplification success in Palaeolithic equid remains from the Alpine foreland. Archaeol Anthropol Sci. 2015;7: 505–515. doi:10.1007/s12520-014-0213-4

5. Burger J, Hummel S, Hermann B, Henke W. DNA preservation: a microsatellite-DNA study on ancient skeletal remains. Electrophoresis. Wiley Subscription Services, Inc., A Wiley Company; 1999;20: 1722–8. doi:10.1002/(SICI)1522-2683(19990101)20:8<1722::AID-ELPS1722>3.0.CO;2-4

6. Swift MJ, Heal OW, Anderson JM. Decomposition in Terrestrial Ecosystems. University of California Press; 1979.

7. Collins MJ, Nielsen-Marsh CM, Hiller J, Smith CI, Roberts JP, Prigodich R V., et al. The survival of organic matter in bone: a review. Archaeometry. 2002;44: 383–394. doi:10.1111/1475-4754.t01-1-00071

8. Alonso a, Andelinović S, Martín P, Sutlović D, Erceg I, Huffine E, et al. DNA typing from skeletal remains: evaluation of multiplex and megaplex STR systems on DNA isolated from bone and teeth samples. Croat Med J. 2001;42: 260–266. doi:10.1016/j.seppur.2013.09.009

9. Leney MD. Sampling skeletal remains for ancient DNA (aDNA): A measure of success. Hist Archaeol. 2006;40: 31–49.

10. Milos A, Selmanović A, Smajlović L, Huel RLM, Katzmarzyk C, Rizvić A, et al. Success rates of nuclear short tandem repeat typing from different skeletal elements. Croat Med J. 2007;48: 486–493.

11. Olszta MJ, Cheng X, Jee SS, Kumar R, Kim Y-Y, Kaufman MJ, et al. Bone structure and formation: A new perspective. Mater Sci Eng R. 2007; doi:10.1016/j.mser.2007.05.001

12. Child AM. Microbial Taphonomy of Archaeological Bone [Internet]. Studies in Conservation. 1995 Feb. doi:10.2307/1506608

13. Schultz M. Microscopic Structure of Bone. Forensic taphonomy: the postmortem fate of human remains. 1997. pp. 187–199.

14. Keenan SW, Summers AS. Early diagenesis and recrystallization of bone. Geochim Cosmochim Acta. 2017;196: 209–223. doi:10.1016/j.gca.2016.09.033

15. Gotherstrom A, Collins MJ, Angerbjorn A, Liden K. Bone preservation and DNA amplification. Archaeometry. 2002;3: 395–404. doi:10.1111/1475-4754.00072

16. Sosa C, Vispe E, Núñez C, Baeta M, Casalod Y, Bolea M, et al. Association between ancient bone preservation and dna yield: a multidisciplinary approach. Am J Phys Anthropol. 2013;151: 102–9. doi:10.1002/ajpa.22262

17. Campos PF, Craig OE, Turner-Walker G, Peacock E, Willerslev E, Gilbert MTP. DNA in ancient bone – Where is it located and how should we extract it? Ann Anat. Elsevier GmbH.; 2012;194: 7–16. doi:10.1016/j.aanat.2011.07.003

18. Graw M, Weisser HJ, Lutz S. DNA typing of human remains found in damp environments. Forensic Sci Int. 2000;113: 91–95. doi:10.1016/S0379-0738(00)00221-8

19. Misner LM, Halvorson AC, Dreier JL, Ubelaker DH, Foran DR. The correlation between skeletal weathering and dna quality and quantity. J Forensic Sci. 2009;54: 822–828. doi:10.1111/j.1556-4029.2009.01043.x

20. Gilbert MTP, Rudbeck L, Willerslev E, Hansen AJ, Smith C, Penkman KEH, et al. Biochemical and physical correlates of DNA contamination in archaeological human bones and teeth excavated at Matera, Italy. J Archaeol Sci. 2005;32: 785–793. doi:10.1016/j.jas.2004.12.008

21. Iwamura ESM, Oliveira CRGCM, Soares-Vieira JA, Nascimento SAB, Mu??oz DR. A Qualitative Study of Compact Bone Microstructure and Nuclear Short Tandem Repeat Obtained From Femur of Human Remains Found on the Ground and Exhumed 3 Years After Death. Am J Forensic Med Pathol. 2005;26: 33–44. doi:10.1097/01.paf.0000154116.30837.d5

22. Andronowski JM, Mundorff AZ, Pratt I V, Davoren JM, Cooper DML. Evaluating differential nuclear DNA yield rates and osteocyte numbers among human bone tissue types: A synchrotron radiation micro-CT approach. 2017;28: 211–218. doi:10.1016/j.fsigen.2017.03.002

23. Haynes S, Searle JB, Bretman A, Dobney KM†. Bone Preservation and Ancient DNA: The Application of Screening Methods for Predicting DNA Survival. J Archaeol Sci. 2002;29: 585–592. doi:10.1006/jasc.2001.0731

24. Jans MMEME, Nielsen-marsh CMM, Smith CII, Collins MJJ, Kars H. Characterisation of microbial attack on archaeological bone. J Archaeol Sci. Academic Press; 2004;31: 87–95. doi:10.1016/j.jas.2003.07.007

25. Edson SM, Christensen AF, Barritt SM, Meehan A, Leney MD, Finelli LN. Sampling of the cranium for mitochondrial DNA analysis of human skeletal remains. Forensic Sci Int Genet Suppl Ser. 2009;2: 269–270. doi:10.1016/j.fsigss.2009.09.029

26. Mundorff AZ, Bartelink EJ, Mar-Cash E. DNA preservation in skeletal elements from the world trade center disaster: Recommendations for Mass Fatality Management. J Forensic Sci. 2009;54: 739–745. doi:10.1111/j.1556-4029.2009.01045.x

27. Hines D, Vennemeyer M, Amory S, Huel R, Hanson I, Katzmarzyk C, et al. Prioritized Sampling of Bone and Teeth for DNA Analysis in Commingled Cases. Commingled Human Remains: Methods in Recovery, Analysis, and Identification. Elsevier Inc; 2014. pp. 275–305.

28. Mundorff A, Davoren JM. Examination of DNA yield rates for different skeletal elements at increasing post mortem intervals. Forensic Sci Int Genet. 2014;8: 55–63. doi:10.1016/j.fsigen.2013.08.001

29. Jans MME, Kars H, Nielsen-Marsh CM, Smith CI, Nord AG, Arthur P, et al. IN SITU PRESERVATION OF ARCHAEOLOGICAL BONE: A HISTOLOGICAL STUDY WITHIN A MULTIDISCIPLINARY APPROACH. Archaeometry. 2002;44: 343–352.

30. Hackett CJ. Microscopical Focal Destruction (Tunnels) in Exhumed Human Bones. Med Sci Law. SAGE PublicationsSage UK: London, England; 1981;21: 243–265. doi:10.1177/002580248102100403

31. White L, Booth TJ. The origin of bacteria responsible for bioerosion to the internal bone microstructure: Results from experimentally-deposited pig carcasses. Forensic Sci Int. 2014;239: 92–102. doi:10.1016/j.forsciint.2014.03.024

32. Booth TJ. An Investigation Into the Relationship Between Funerary Treatment and Bacterial Bioerosion in European Archaeological Human Bone. Archaeometry. 2016;58: 484–499. doi:10.1111/arcm.12190

33. Balzer A, Gleixner G, Grupe G, Schmidt H-L, Schramm S, Turban-Just S. In vitro decomposition of bone collagen by soil bacteria: the implications for stable isotope analysis in archaeometry. Archaeometry. 1997;39: 415–429. doi:10.1111/j.1475-4754.1997.tb00817.x

34. Child AMM. Towards an Understanding of the Microbial Decomposition of Archaeological Bone in the Burial Environment. J Archaeol Sci. Academic Press; 1995;22: 165–174. doi:10.1006/JASC.1995.0018

35. Marchiafava V, Bonucci E, Ascenzi A. Fungal Osteoclasia: a Model of Dead Bone Resorption. Calcif Tissue Res. Springer-Verlag; 1974;14: 195–210. Available: https://link-springer-com.proxy.lib.utk.edu/content/pdf/10.1007%2FBF02060295.pdf

36. Staley JT, Konopka A. MEASUREMENT OF IN SITU ACTI\TITIES OF NONPHOTOSYNTHETIC MICROORGANISMS IN AQUATIC AND TERRESTRIAL HABITATS. Annu Rev Microbiol. 1985;39: 321–346. Available: www.annualreviews.org

37. Damann FE, Williams DE, Layton AC. Potential Use of Bacterial Community Succession in Decaying Human Bone for Estimating Postmortem Interval · ·. J Forensic Sci. 2015;60: 844–850. doi:10.1111/1556-4029.12744

38. Reeb V, Kolel A, McDermott TR, Bhattacharya D. Good to the bone: Microbial community thrives within bone cavities of a bison carcass at Yellowstone National Park. Environ Microbiol. 2011;13: 2403–2415. doi:10.1111/j.1462-2920.2010.02359.x

39. Amory S, Huel R, Bilić A, Loreille O, Parsons TJ. Automatable full demineralization DNA extraction procedure from degraded skeletal remains. Forensic Sci Int Genet. 2012;6: 398–406. doi:10.1016/j.fsigen.2011.08.004

40. Castillo M, Martín-Orúe SM, Manzanilla EG, Badiola I, Martín M, Gasa J. Quantification of total bacteria, enterobacteria and lactobacilli populations in pig digesta by real-time PCR. Vet Microbiol. 2006;114: 165–170. doi:10.1016/j.vetmic.2005.11.055

41. Klindworth A, Pruesse E, Schweer T, Peplies J, Quast C, Horn M, et al. Evaluation of general 16S ribosomal RNA gene PCR primers for classical and next-generation sequencing-based diversity studies. Nucleic Acids Res. Oxford University Press; 2013;41: e1. doi:10.1093/nar/gks808

42. Hugerth LW, Muller EEL, Hu YOO, Lebrun LAM, Roume H, Lundin D, et al. Systematic design of 18S rRNA gene primers for determining eukaryotic diversity in microbial consortia. PLoS One. 2014;9. doi: 10.1371/journal.pone.0095567

43. Martin M. Cutadapt removes adapter sequences from high-throughput sequencing reads kenkyuhi hojokin gan rinsho kenkyu jigyo. EMBnet.journal. 2013;17: 10–12. doi:10.14806/ej.17.1.200

44. Bolger AM, Lohse M, Usadel B. Trimmomatic: A flexible read trimming tool for Illumina NGS data. Bioinformatics. 2014;btu170.

45. Schloss PD, Westcott SL, Ryabin T, Hall JR, Hartmann M, Hollister EB, et al. Introducing mothur: Open-source, platform-independent, community-supported software for describing and comparing microbial communities. Appl Environ Microbiol. 2009;75: 7537–7541. doi:10.1128/AEM.01541-09

46. R Core Team. R: A language and environment for statistical computing. Foundation for Statistical Computing, Vienna, Austria.; 2018.

47. McMurdie PJ, Holmes S. Phyloseq: a bioconductor package for handling and analysis of high-throughput phylogenetic sequence data. Pac Symp Biocomput. NIH Public Access; 2012; 235–46. Available: http://www.ncbi.nlm.nih.gov/pubmed/22174279

48. Bradley IM, Pinto AJ, Guest JS. Design and Evaluation of Illumina MiSeq-Compatible, 18S rRNA Gene-Specific Primers for Improved Characterization of Mixed Phototrophic Communities. Appl Environ Microbiol. American Society for Microbiology; 2016;82: 5878–91. doi:10.1128/AEM.01630-16

49. Wuertz D, Setz T, Chalabi Y. fBasics: Rmetrics – Markets and Basic Statistics [Internet]. Available: https://cran.r-project.org/package=fBasics

50. Fox J, Weisberg S. An {R} Companion to Applied Regression [Internet]. 2nd Editio. Thousand Oaks CA: Sage; 2011. Available: http://socserv.socsci.mcmaster.ca/jfox/Books/Companion

51. Lüdecke D. _sjstats: Statistical Functions for Regression Models [Internet]. 2019. doi:10.5281/zenodo.1284472

52. Steinberger A. seq-scripts release v. 1.0. 2016. doi:10.5281/zenodo.1458243

53. Pedregosa F, Varoquax G, Gramfort A, Michel V, Grisel O, Blondel M, et al. Scikit-learn: Machine Learning in Python. Matthieu Perrot [Internet]. Journal of Machine Learning Research. 2011. Available: http://scikit-learn.sourceforge.net.

54. Cobaugh KL, Schaeffer SM, DeBruyn JM. Functional and Structural Succession of Soil Microbial Communities below Decomposing Human Cadavers. PLoS One. 2015;10: e0130201. doi:10.1371/journal.pone.0130201

55. Adserias-Garriga J, Hernández M, Quijada NM, Lázaro DR, Steadman D, Garcia-Gil J. Daily thanatomicrobiome changes in soil as an approach of postmortem interval estimation: An ecological perspective. Forensic Sci Int. 2017;278: 388–395. doi:10.1016/j.forsciint.2017.07.017

56. Carter DO, Tibbett M. Taphonomic mycota: fungi with forensic potential. J Forensic Sci. ASTM International; 2003;48: 168–171. doi:JFS2002169_481

57. Dumack K, Flues S, Hermanns K, Bonkowski M. Rhogostomidae (Cercozoa) from soils, roots and plant leaves (Arabidopsis thaliana): Description of Rhogostoma epiphylla sp. nov. and R. cylindrica sp. nov. Eur J Protistol. 2017;60: 76–86. doi:10.1016/j.ejop.2017.06.001

58. Metcalf JL, Wegener Parfrey L, Gonzalez A, Lauber CL, Knights D, Ackermann G, et al. A microbial clock provides an accurate estimate of the postmortem interval in a mouse model system. Elife. 2013;2: e01104. doi:10.7554/eLife.01104

59. Keenan SW, Emmons AL, Taylor LS, Phillips G, Mason AR, Mundorff AZ, et al. Spatial impacts of a multi-individual grave on microbial and microfaunal communities and soil biogeochemistry. Yang Z, editor. PLoS One. Public Library of Science; 2018;13: e0208845. doi:10.1371/journal.pone.0208845

60. Szelecz I, Sorge F, Seppey CVW, Mulot M, Steel H, Neilson R, et al. Effects of decomposing cadavers on soil nematode communities over a one-year period. Soil Biol Biochem. 2016;103: 405–416. doi:10.1016/j.soilbio.2016.09.011

61. Szelecz I, Losch, Sandra, Seppey, Christophe V.W., Lara, Enrique, Singer, David, Sorge, Franziska, et al. Comparative analysis of bones, mites, soil chemistry, nematodes and soil microeukaryotes from a suspected homicide to estimate the post-mortem interval. Nat Sci Reports. 2018;8: 1:14.

62. Szelecz I, Fournier B, Seppey C, Amendt J, Mitchell E. Can soil testate amoebae be used for estimating the time since death? A field experiment in a deciduous forest. Forensic Sci Int. 2014;236: 90–98. doi:10.1016/j.forsciint.2013.12.030

63. Seppey CVW, Fournier B, Szelecz I, Singer D, Mitchell EAD, Lara E. Response of forest soil euglyphid testate amoebae (Rhizaria: Cercozoa) to pig cadavers assessed by high-throughput sequencing. Int J Legal Med. 2016;130: 551–562. doi:10.1007/s00414-015-1149-7

64. Theodorakopoulos N, Bachar D, Christen R, Alain K, Chapon V. Exploration of Deinococcus-Thermus molecular diversity by novel group-specific PCR primers. Microbiologyopen. 2013;2: 862–872. doi:10.1002/mbo3.119

65. Wilson-Taylor RJ, Dautartas AM. Time Since Death Estimation and Bone Weathering. In: Langley NR, Tersigni-Tarrant MA, editors. Forensic Anthropology A Comprehensive Introduction. second edi. 2017.

66. Fierer N, Hamady M, Lauber CL, Knight R. The influence of sex, handedness, and washing on the diversity of hand surface bacteria. Proc Natl Acad Sci. National Academy of Sciences; 2008;105: 17994–17999. doi:10.1073/PNAS.0807920105

67. De Filippo C, Cavalieri D, Di Paola M, Ramazzotti M, Poullet JB, Massart S, et al. Impact of diet in shaping gut microbiota revealed by a comparative study in children from Europe and rural Africa. Proc Natl Acad Sci U S A. National Academy of Sciences; 2010;107: 14691–6. doi:10.1073/pnas.1005963107

68. HMP Consortium THMP, Huttenhower C, Gevers D, Knight R, Abubucker S, Badger JH, et al. Structure, function and diversity of the healthy human microbiome. Nature. Nature Publishing Group; 2012;486: 207–214. doi:10.1038/nature11234

69. Larsen N, Vogensen FK, van den Berg FWJ, Nielsen DS, Andreasen AS, Pedersen BK, et al. Gut Microbiota in Human Adults with Type 2 Diabetes Differs from Non-Diabetic Adults. Bereswill S, editor. PLoS One. Public Library of Science; 2010;5: e9085. doi:10.1371/journal.pone.0009085

70. Pechal JL, Schmidt CJ, Jordan HR, Benbow ME. A large-scale survey of the postmortem human microbiome, and its potential to provide insight into the living health condition. Nat Sci Reports. 2018;8. doi: 10.1038/s41598-018-23989-w

71. Caporaso JG, Lauber CL, Costello EK, Berg-Lyons D, Gonzalez A, Stombaugh J, et al. Moving pictures of the human microbiome. Genome Biol. BioMed Central; 2011;12: R50. doi: 10.1186/gb-2011-12-5-r50

72. Costello EEK, Lauber CCL, Hamady M, Fierer N, Gordon JI, Knight R. Bacterial community variation in human body habitats across space and time. Science (80-). 2009;326: 1694–7. doi:10.1126/science.1177486.Bacterial

73. Fierer N, Lauber CL, Zhou N, McDonald D, Costello EK, Knight R. Forensic identification using skin bacterial communities. Proc Natl Acad Sci U S A. 2010;107: 6477–6481. doi:10.1073/pnas.1000162107

74. Lax S, Hampton-Marcell JT, Gibbons SM, Colares GB, Smith D, Eisen JA, et al. Forensic analysis of the microbiome of phones and shoes. Microbiome. BioMed Central; 2015;3: 21. doi:10.1186/s40168-015-0082-9

75. Nicholson RA. Bone Degradation, Burial Medium and Species Representation: Debunking the Myths, an Experiment-based Approach [Internet]. Journal of Archaeological Science; 1996;23: 513–533.

76. Metcalf JL, Xu ZZ, Weiss S, Lax S, Treuren W Van, Hyde ER, et al. Microbial community assembly and metabolic function during mammalian corpse decomposition. Science (80-). 2016;351: 158–162. doi:10.1126/science.aad2646

77. Ghannoum MA, Jurevic RJ, Mukherjee PK, Cui F, Sikaroodi M, Naqvi A, et al. Characterization of the Oral Fungal Microbiome (Mycobiome) in Healthy Individuals. May RC, editor. PLoS Pathog. Public Library of Science; 2010;6: e1000713. doi:10.1371/journal.ppat.1000713

78. Nielsen-Marsh CM, Hedges REM. Patterns of Diagenesis in Bone I: The Effects of Site Environments. J Archaeol Sci. 2000;27: 1139–1150. doi:10.1006/jasc.1999.0537

79. Trueman CN, Martill DM. The long-term survival of bone: the role of bioerosion. Archaeometry. Wiley/Blackwell (10.1111); 2002;44: 371–382. doi:10.1111/1475-4754.t01-1-00070

80. Hollund HI, Teasdale MD, Mattiangeli V, Sverrisdóttir OÓ, Bradley DG, O’connor T. Pick the Right Pocket. Sub-sampling of Bone Sections to Investigate Diagenesis and DNA Preservation. Int J Osteoarchaeol. 2017;27: 365–374. doi:10.1002/oa.2544

